# Wide distribution of alternatively coded Lak megaphages in animal microbiomes

**DOI:** 10.1101/2021.01.08.425732

**Authors:** Marco A. Crisci, Lin-Xing Chen, Audra E. Devoto, Adair L. Borges, Nicola Bordin, Rohan Sachdeva, Adrian Tett, Allison M. Sharrar, Nicola Segata, Francesco Debenedetti, Mick Bailey, Rachel Burt, Rhiannon M. Wood, Lewis J. Rowden, Paula M. Corsini, Mark A. Holmes, Shufei Lei, Jillian F. Banfield, Joanne M. Santini

## Abstract

Lak phages with alternatively coded ~540 kbp genomes were recently reported to replicate in *Prevotella* in the gut microbiomes of humans that consume a non-western diet, baboons and some pigs. Here, we investigate the diversity and broader distribution of Lak phages in human and animal microbiomes using diagnostic PCR and genome-resolved metagenomics. Lak phages were detected in 13 different animal types and are particularly prevalent in pigs, with significant enrichment in the hindgut compared to foregut. We reconstructed 34 new Lak genomes, including six curated complete genomes, all of which are alternatively coded. The most deeply branched Lak is from a horse faecal sample and is the largest phage genome from an animal microbiome (~660 kbp). From the Lak genomes, we identified families of hypothetical proteins associated with specific animal types. Overall, we substantially expanded Lak phage diversity and demonstrate their occurrence in a variety of human and animal microbiomes.

## Introduction

*Prevotella* and *Bacteroides* (phylum *Bacteroidetes*) occupy similar ecological niches and compete for resources in gut microbiomes ^1,2^. *Prevotella* and *Bacteroides*-dominated enterotypes are linked to non-western and western diets, respectively ^3–6^. Diets low in fat and protein but high in fibre promote *Prevotella* growth, whereas diets high in animal fat, protein and starch promote *Bacteroides* growth ^2,4^. *Prevotella* can metabolise fibre and produce volatile fatty acids that are crucial to gut health more effectively than *Bacteroides* ^7^. *Prevotella* are also widespread in pig gut microbiomes and are generally associated with improved growth performance, an observation of interest because pigs are important production animals and model for the human gut ^8–10^.

Lak megaphages that replicate in *Prevotella* were recently discovered in human and baboon gut microbiomes using genome-resolved metagenomics ^11^. To date, these phages have the largest genomes identified in gut microbiomes (> 540 kbp in length). Lak phage sequences were also detected in Danish pig metagenomes abundant in *Prevotella*, and in cow rumens at low abundance ^11^. Unlike smaller *Prevotella* phages that typically adopt a temperate lifestyle ^12,13^, Lak genomes do not contain identifiable integrases and no prophages have been detected in bacterial chromosomes. Thus, it is likely that Lak phages have a lytic life cycle. Lysis by Lak phages could alter the composition and abundance of *Prevotella* in the animal/human host, affecting microbial community structure and nutrient availability.

A notable feature of Lak is the use of an alternative genetic code, where the “TAG” stop codon is repurposed to encode glutamine (Q) ^11^. Lak genomes encode a suppressor tRNA with a CTA anticodon needed to repurpose TAG. Moreover, presence of release factor 2 (RF2) terminates protein translation through recognition of TGA and TAA stop codons but not TAG. The reason for Lak phage codon reassignment is unknown, but it may disrupt translation of bacterial genes^14^.

In this study, we screened digesta/faecal and mucosal samples from animals that consume dietary fibre by PCR, revealing the distribution of Lak phages in gut microbiomes that may harbour *Prevotella*. We also quantified the abundance of Lak phage and *Prevotella* across the swine gastrointestinal tract (GIT) and vagina by qPCR. Metagenomic datasets from new and previously sequenced samples were investigated to define Lak phage diversity, gene content and genetic code. From these metagenomes, we manually curated six new Lak genomes to completion, substantially expanding the genome size range and uncovering the largest phage genome reported from an animal microbiome. In addition to expanding clade diversity via sampling of 34 new Lak genomes, bacterial hosts and evolutionary relationships were predicted, and the extent and origins of alternative codon usage among Lak phages evaluated. Protein family analyses were performed to identify animal-specific protein clusters that may be important for adaptation of Lak phages to their microbiome environments.

## Results

### Lak phages detected in various animal gut samples by PCR

PCR primer sets targeting the major capsid protein (MCP), tail sheath monomer (TSM) and portal vertex protein (PVP) signature genes detected Lak in faecal and mucosal samples from many animal microbiomes (Table 1, Supplementary Fig. S1, Supplementary Tables 1 and 2). Lak was detected in 80% of pigs (*n* = 28), but was undetectable in gestating sows (*n* = 4) and a post-farrow sow (*n* = 1) with piglets (*n* = 2). Jejunal and ileal (foregut), and proximal spiral, distal spiral, caecal and rectal (hindgut) lumen and mucosal samples from six Bristol finisher pigs tested positive and were subjected to qPCR quantification. Three of five vaginal samples tested positive from pigs where Lak was detected in the rectum, but not in the lungs, although *Prevotella* 16S rRNA genes were detected at all body sites by PCR. A subset of PCR products from each animal cohort were sequenced, confirming the presence of Lak (Table 1; Supplementary Table 1).

**Table 1.** Animal gut samples tested positive by PCR diagnostics. A subset of PCR products from each cohort were sequenced to confirm the presence of Lak. ^1^ Post-mortem pig samples from: foregut (jejunum and ileum) and hindgut (caecum, proximal spiral, distal spiral and rectum). ^2^ Lak was detected in all rectums (n=5) and 3/5 vaginal mucosa, but not in lungs of the same animals. See details in Supplementary Table 2.

| Animal | Sample Type | Details | PCR positive |
| --- | --- | --- | --- |
| Cow (*Bos taurus*), Holstein | Rumen fluid | 1 Individual, female | 1 / 3 |
| Warthog (*Phacochoerus africanus*) | Faeces | Group of 2, pooled | 1 / 3 |
| White-naped mangabey (*Cercocebus lunulatus*) | Faeces | Group of 7, pooled | 1 / 3 |
| Galapagos giant tortoise (*Chelonoidis nigra*) | Faeces | Adult group of 3 and juvenile group of 3, pooled separately | 2 / 2 |
| Fallow Deer (*Dama dama*) | Faeces | 1 Individual, wild | 1 / 1 |
| Horse (*Equus ferus caballus*), racing thoroughbred | Faeces | 6 Individual over 3 days, various sexes and ages | 12 / 18 |
| (Cambridge) Pigs (*Sus scrofa*), Large White/Landrace/Hampshire | Faeces | 19 Individual, various sexes and ages | 12 / 19 |
| (Bristol) Pigs (*Sus scrofa*), Welsh and Welsh/Pettrain | Lumen digesta and mucosal scrapings <sup>1</sup> | 6 Individual, finisher pigs, various sexes. | 70 / 70 |
| (Bristol) Pigs (*Sus scrofa*), Welsh/Pettrain | Vaginal and Rectal Samples <sup>2</sup> | 5 Individual, finisher pigs, female | 8 / 15 |
| (Royal Veterinary College) Pigs (*Sus scrofa*), Large White / Unknown cross | Faeces | Various groups, pooled | 5 / 5 |

### Lak phage abundance across the pig gastrointestinal tract and vagina revealed by qPCR

The abundance of Lak and *Prevotella* was quantified in triplicate across the GIT of the Bristol finisher pigs (Table 1, Fig. 1a, Supplementary Figs. S2-5; Methods). Lak abundance correlated with *Prevotella* abundance across the entire GIT, although there were fewer Lak phage than *Prevotella* copies at all sites (Fig. 1b). Together, GIT site and sample type (mucosa or lumen) had a significant effect on *Prevotella* (*P* = 0.019) and Lak phage (*P* = 0.003) abundance. Foregut Lak and *Prevotella* abundance (jejunum and ileum) were significantly lower than hindgut sites (caecum, proximal spiral, distal spiral) in both the lumen and mucosa (*P* < 0.01; Tukey’s HSD; Fig. 1b). However, there was no statistically significant difference in both Lak and *Prevotella* abundance between the lumen and mucosa at each GIT site (*P* > 0.05; Tukey’s HSD; Fig. 1b). The ratio of Lak: *Prevotella* copy numbers did not differ between mucosa and lumen at each site, but generally, Lak phage: *Prevotella* copy numbers were significantly higher in the foregut mucosa (jejunum = 0.076; Ileum = 0.041) than in hindgut lumen and mucosal sites (range = 0.0002-0.003; *P* < 0.05; Tukey’s HSD; Fig. 1c). The proportion of Lak: *Prevotella* in the lumen was also higher in the foregut than in hindgut lumens, except in the caecum. Within the all-female pig group, Lak phages (*P* = 0.0001) and *Prevotella* (*P* = 0.0002) were significantly less abundant in the vagina than in the rectum (Supplementary Table 3).

**Fig. 1.**
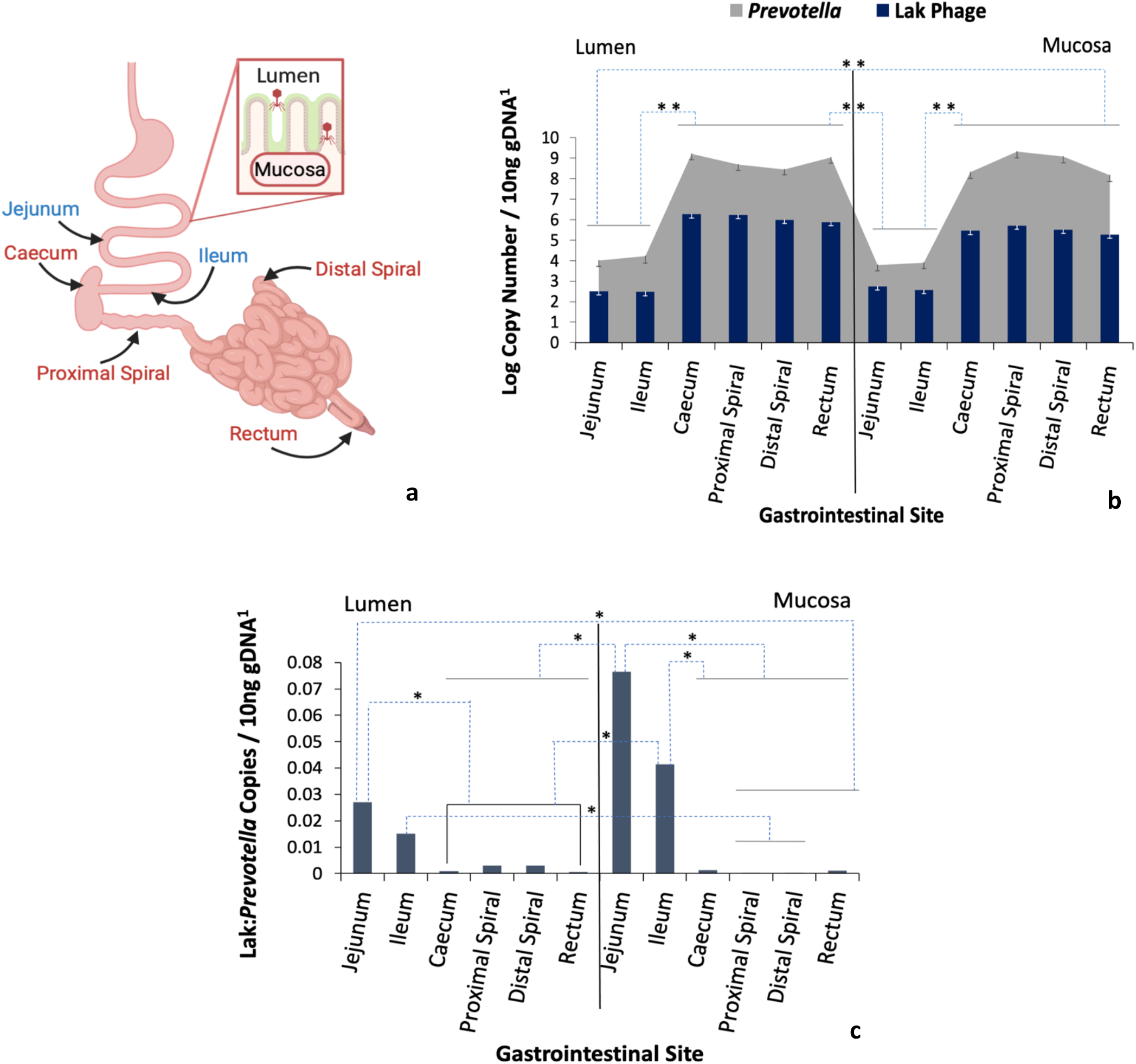
Abundance of Lak phage and *Prevotella* in the pig GI tract. (a) Schematic of pig GI tract with labels indicating the sites sampled: Blue labels = foregut, Red labels = hindgut (main sites of microbial fibre fermentation) (created using <u>BioRender</u>). (b) Difference in Lak phage log mean abundance across pig lumen and mucosal sites coincides with *Prevotella* abundance. Error bars show standard error of the least squared mean. (c) Ratios of Lak phage to *Prevotella* genome copies. Anti-log means are presented but log values were analysed, ranging from −1.12 to −3.67 with standard errors of 0.28-0.35. For both (b) and (c), ^1^ Lak phage and total *Prevotella* copy number determined by absolute quantification qPCR using the standard curve method, with 10 ng pooled DNA from each GI site from 6 pigs. Lak primers targeted the major capsid gene, and *Prevotella* primers targeted the 16S rRNA gene. Blue dotted lines represent differences in abundances deemed significantly significant by Tukey’s HSD test: \*\**P* < 0.001 and \**P* < 0.05. All other comparisons lacked statistical significance (*P* > 0.05). Black lines represent grouped GIT sites that did not differ from each other (*P* > 0.05), but differed from other sites. See details in Supplementary Fig. S5.

### Newly reconstructed Lak phage genomes

Eight novel Lak phage genomes were reconstructed from new metagenomic datasets for a subset of samples identified to contain Lak, and 28 were reconstructed from published metagenomic datasets. Of these, 8 came from pig faecal samples (including one from a published dataset of Danish pig, Pig_ID_1901_F52), 18 from human fecal samples, 7 genomes from baboon faecal samples and one from a horse faecal sample (Table 2). Six of these 34 draft genomes were manually curated to completion. Two genomes are ~476 kbp in length (476,085 and 476,118 bp, representative of a set of 4 genomes with GC contents of ~31% from pig gut microbiomes), one genome is 517,629 bp in length (GC content ~26% from a pig, detectable by read mapping at ≥ 5X coverage in 38.4% of previously reported pig metagenomes ^15^) and one 659,950 bp genome (GC content ~ 29% from a horse faecal sample; Table 2, Supplementary Table 1). These findings substantially expand the known range of genome sizes for Lak phages. The ~518 kbp, ~476 kbp (GC31) and ~540 kbp (GC26) genomes are syntenic, and small blocks of sequence account for the differences in genome lengths. However, the 660 kbp phage genome is too divergent at the nucleotide level to align with those of the other clades, and its classification as Lak is based on phylogeny of Lak proteins.

**Table 2.**
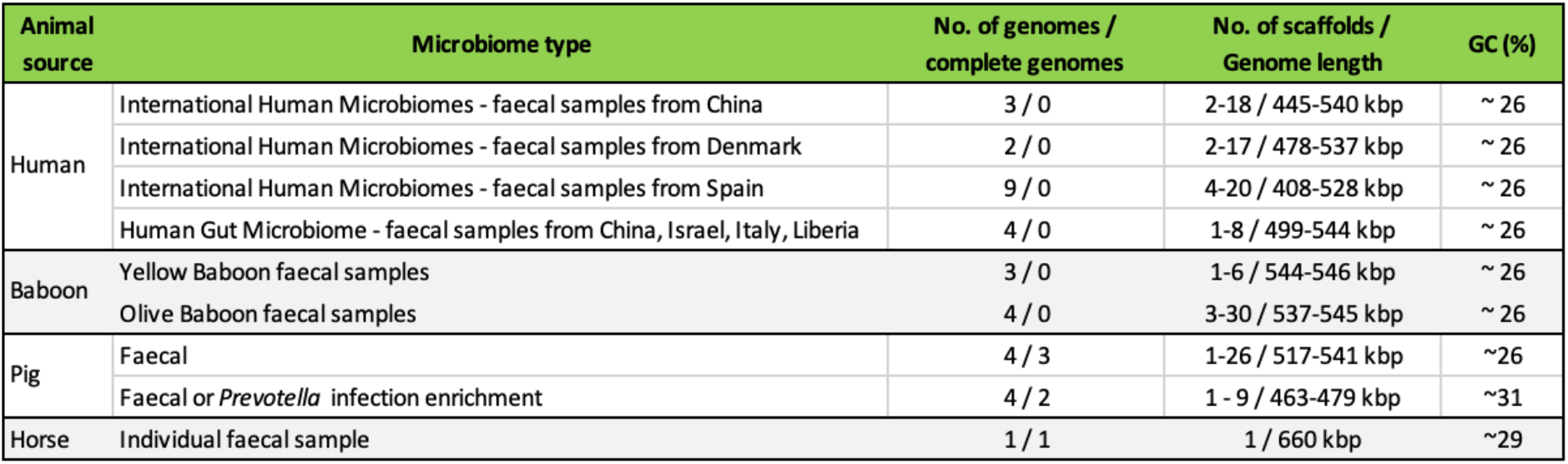
General features of the 34 new Lak phage genomes reconstructed in this study. All the new Lak genomes were included for protein family analyses, along with the 15 published Lak genomes ^11^ and all the 181 circular huge phage genomes reported recently ^16^.

### Phylogenetic relatedness of Lak phages

To investigate the relatedness of Lak phages, phylogenetic trees were constructed based on the PCR-amplified and genome-derived nucleotide sequences of Lak MCP (Fig. 2), TSM and PVP genes (Supplementary Figs. 6 and 7). With all conserved genes, we found that the Lak phages from olive baboon, mangabey, guenons, western red colobus and yellow baboon were more phylogenetically related. The Lak phages from horse, warthog, giant tortoise, cow, fallow deer and most pig microbiomes were generally clustered together on the trees. Moreover, the Lak phages detected in crab-eating macaques were closely related to some from human microbiomes (Fig. 2).

**Fig. 2.**
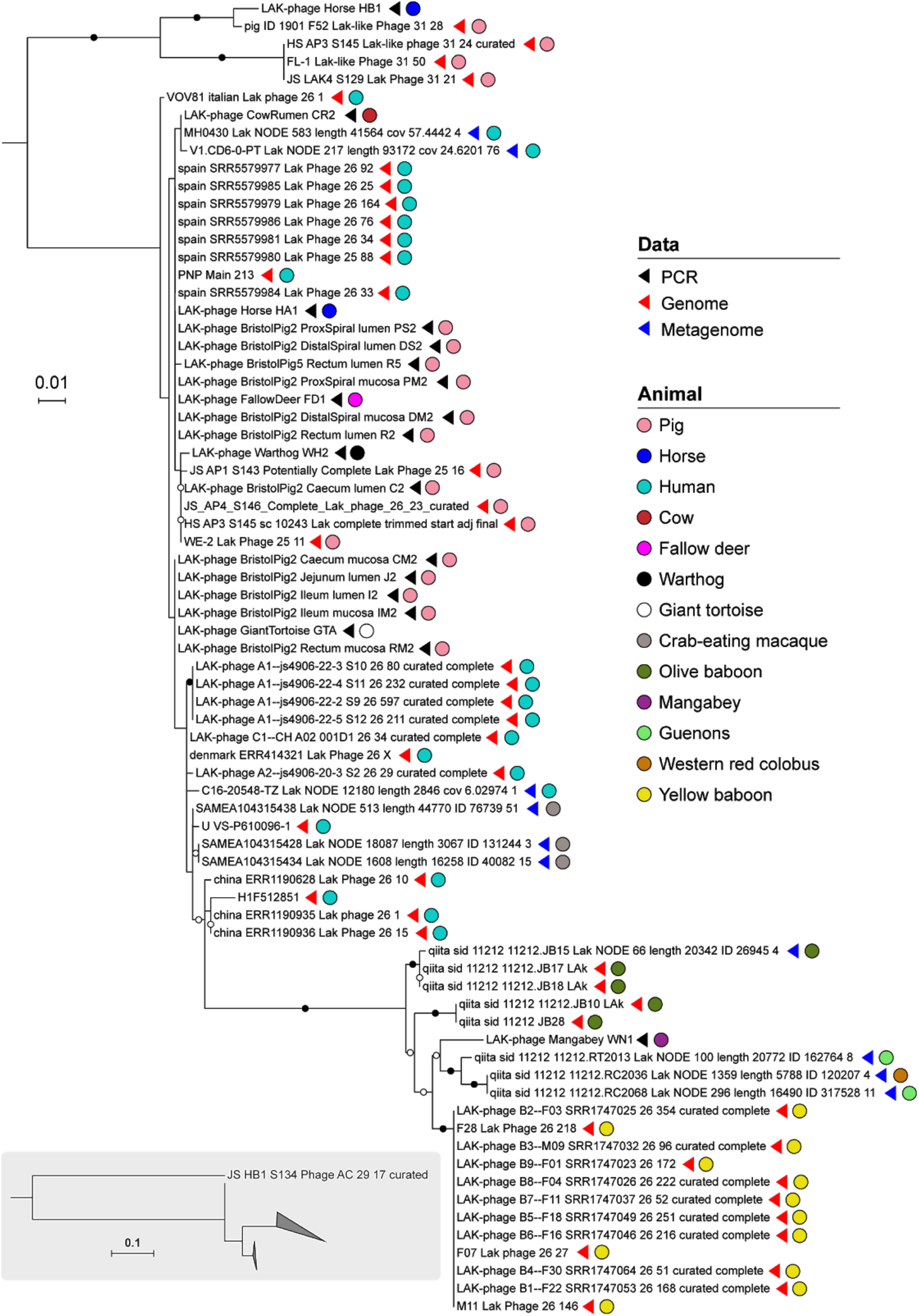
Phylogenetic analyses of Lak based on the sequences from PCR, genomes and metagenomes. The nucleotide sequences encoding the major capsid protein were aligned and trimmed so that all lengths corresponded with that of the PCR sequences. The capsid of the ~660 kbp phage is very divergent from others, thus was excluded from the tree to enable resolution of the other sequences. The phylogenetic relatedness of the ~660 kbp and other Lak phages is shown in the inserted subfigure. Bristol pig sequences obtained from the vaginal mucosa were identical to those found in the digestive tract. Corresponding trees for portal vertex and tail sheath monomer genes are shown in Supplementary Figs. 6 and 7.

### Predicted host(s) of Lak phages

We analysed all of the detected CRISPR-Cas systems from the scaffolds of the corresponding samples. For a given scaffold with a CRISPR-Cas system identified, all spacers from the scaffold and also the reads that mapped to it were extracted to search for their targets (≥90% identity; Methods). We found that the pig-derived WE-2_Lak_Phage_25_11 was targeted by three spacers (total count = 11) from WE-2_scaffold_6241 (total count = 89, unique spacers count = 38; Supplementary Fig. S8). The genome of denmark_ERR1305877_Lak_Phage_26_8 was targeted by two unique spacers (total count = 3), which were respectively from two CRISPR-Cas systems on two scaffolds. None of the scaffolds were binned to a genome, but most of the genes on them had the highest similarity to *Prevotella* genes. The indication that *Prevotella* is the host for these newly reported Lak phages is consistent with the previous finding ^11^ that Lak are targeted by CRISPR spacer matches from *Prevotella* in human gut microbiomes. This putative host is currently classified as CAG 386 which is in the species-level “Clade B” of the *P. copri* complex^17^. We also detected no integrases by functional annotation, corroborating previous findings that Lak phages do not integrate into host genomes ^11^.

### Use of code 15 is conserved throughout the expanded Lak clade

Although we anticipate that Lak phage genomes use genetic code 15 (only TGA and TAA are stop codons), we first predicted the Lak phage genes using code 11 (in which TAG, TGA and TAA are read as stop codons) to check the expanded dataset for evidence of alternative coding. For all Lak, the coding density was consistently low when genes were predicted using code 11, indicating a stop codon reassignment. Re-prediction without use of the TAG stop codon (as in code 15) resulted in full length open reading frames. However, even after re-prediction using code 15, some regions still had low coding density (many regions > 1 kbp and some > 2 kbp with no predicted open reading frames), extending prior findings ^11^.

To determine the phylogenetic span of alternative coding in Lak phages, we searched the metagenome datasets for Lak terminase proteins (whether or not they were on genome fragments assigned to bins). The terminase proteins were highly fragmented when TAG was read as a stop codon.

Coding was uncertain for one group of Lak phages, represented by very short genome fragments (pale blue boxes; Fig. 3). However, results generally indicate that alternative coding persisted from the common Lak phage ancestor. The deepest branches in Fig. 3 represent phages that show no evidence of recoding (Supplementary Fig. S9).

**Fig. 3.**
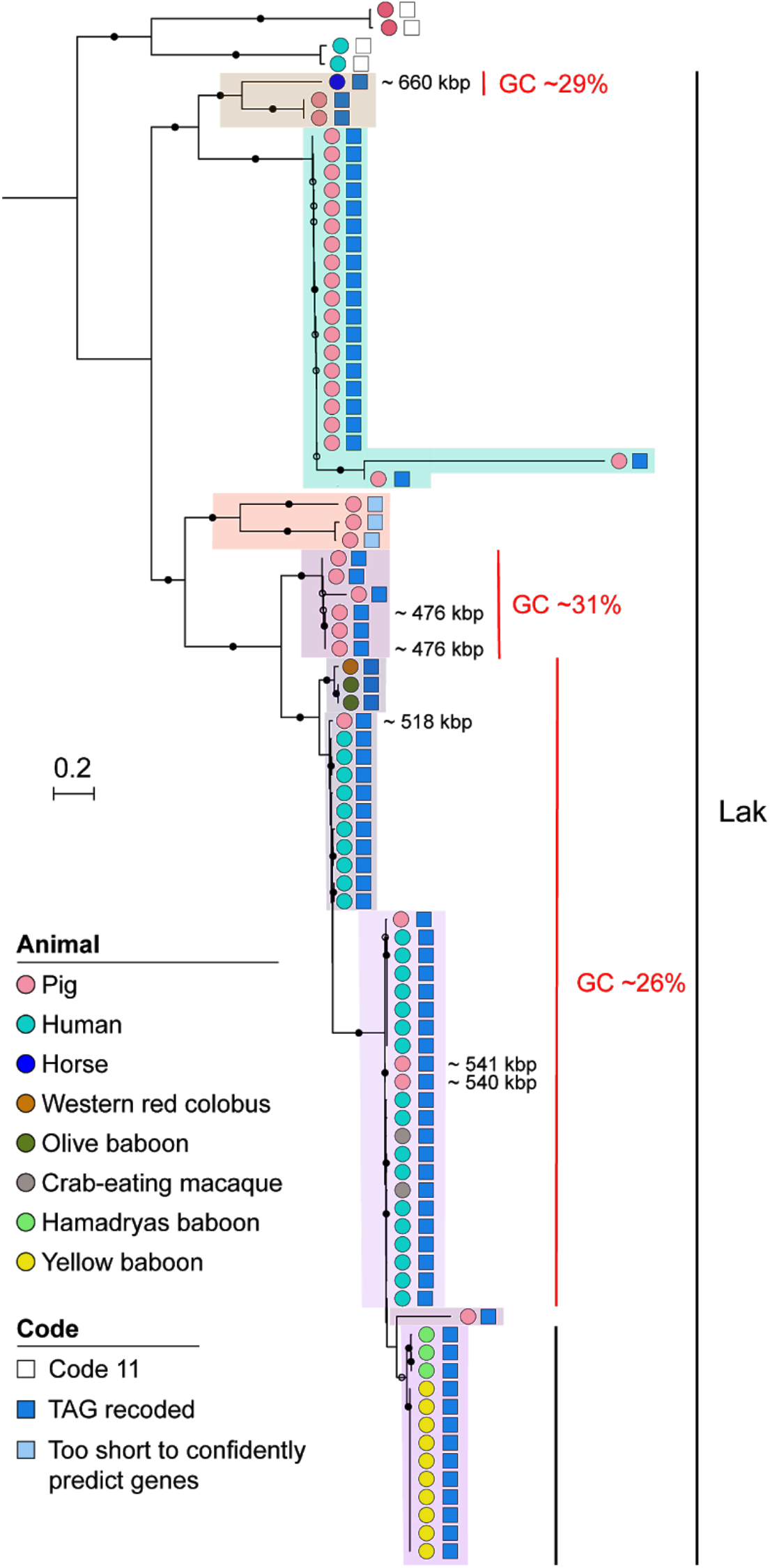
Alternative coding is a persistent feature of Lak phages. The maximum likelihood phylogenetic tree (iqtree (v1.6.12) using the “LG+G4” model (-bb = 1000)) was constructed using sequences for the terminase protein sequence. The genome sizes shown are based on those of complete Lak phages in each clade. Nodes with ≥ 90% bootstrap support values are indicated by filled black circles and nodes with 70-90 % support by open circles. Recoding of the TAG stop codon was detected through the Lak lineages but not in phages represented by the deepest branch.

Previous analyses of Lak ^11^ and other alternatively coded phages ^14^ suggested that TAG is recoded to glutamine (Gln, Q). However, prior studies did not investigate variation in TAG codon use patterns within genes, or consider the possibility of alternative translations. Thus, we aligned terminase sequences where TAG is represented as * to identify the aligned amino acids for each clade (Fig. 3). Based on cases where one specific amino acid in at least two different sequences was aligned with one or more * (Fig. 4), we deduced that throughout much of the Lak clade, TAG is likely translated as Q. These in-frame TAGs were probably introduced by synonymous substitution, i.e., CAG (Q) to TAG. In some cases, * aligned with E (glutamic acid), which is chemically similar to Q (Fig. 4). Plausibly, this occurred by mutation of GAG to TAG. Within the four Lak lineages (shaded in Figs. 3 and 4), positions with only * within and across clades may be mutations that introduced TAG after the rise of alternative coding in the ancestral group. Due to low information content, the alignment could not resolve the translation in three clades (green, orange and brown shading in Figs. 3 and 4).

**Fig. 4.**
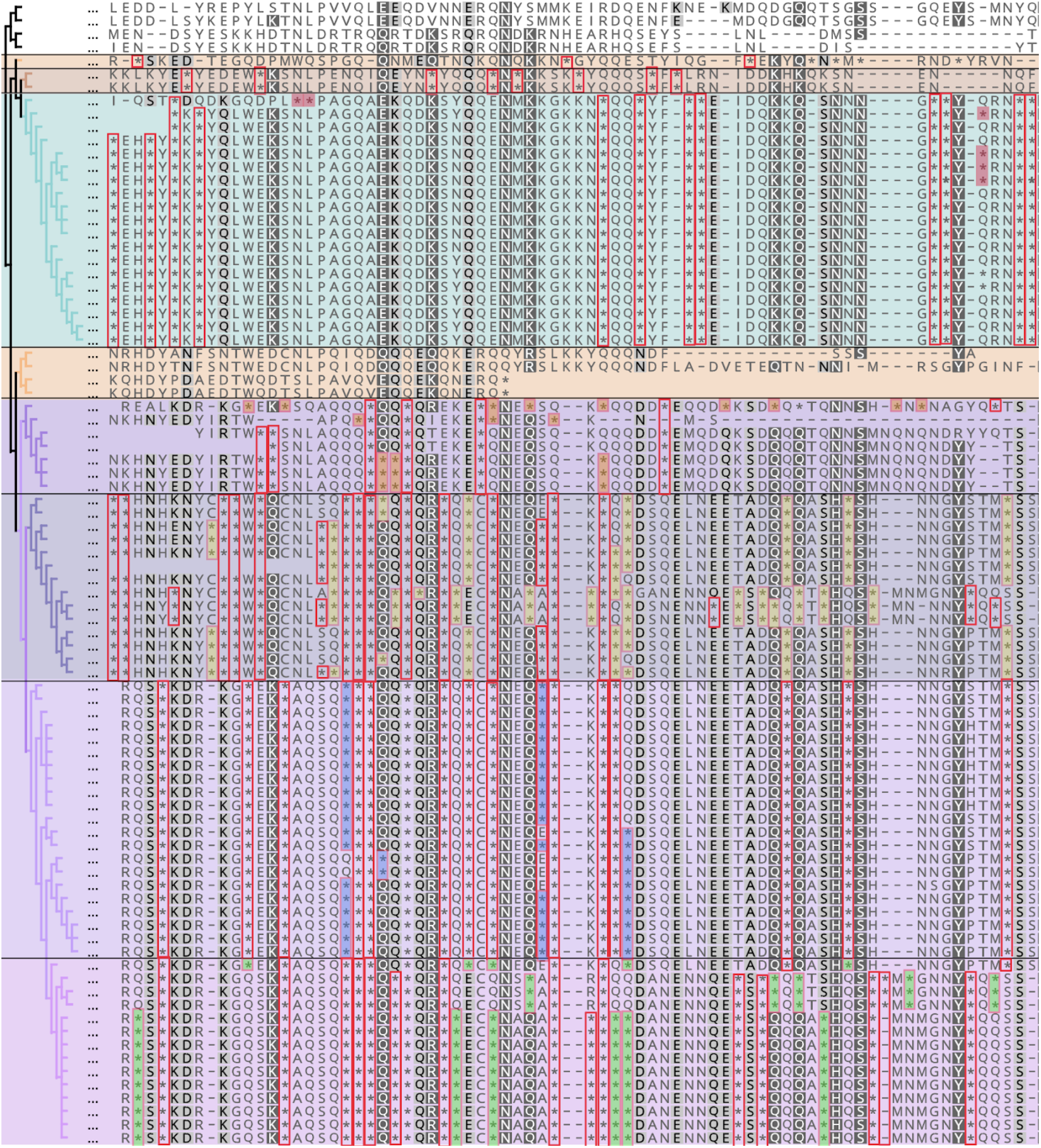
Compressed version of the terminase protein sequence alignment in which all positions except those with in-frame TAG codons (represented by *) have been deleted. Background shading indicates different Lak phage lineages, as shown in Fig. 3). Colors superimposed on * indicate positions in which there is within-clade consensus as to the identity of the aligned amino acid. In the Lak clades with ~26% GC (bottom three groups), Q is the aligned amino acid in 77%, 75% and 85% of cases. There is insufficient information in other groups to predict how TAG is translated.

### tRNA analysis

Stop-codon reassignment can be facilitated by the acquisition of a suppressor tRNA to decode the reassigned stop codon as an amino acid. To define the tRNA repertoire of the expanded Lak clade, we searched the high quality Lak genomes for tRNAs with tRNAscan-SE ^18^. Lak phages encode 24 to 56 tRNAs (Supplementary Table 4), and the majority of phages (36/49) encode 1-2 copies of a suppressor tRNA predicted to decode the TAG stop codon. Notably, these phages also universally (49/49) encode a suppressor tRNA predicted to bind the TAA stop codon. However, we find no other evidence to suggest the TAA stop codon is also recoded in Lak phages.

### Conserved, lineage-specific and animal-specific protein families

We clustered predicted protein sequences into protein families and examined the distribution of families across 49 high quality Lak genomes to investigate whether, and to what extent, Lak phages have a conserved core gene set and if some genes are specific to Lak phages found in gut microbiomes of certain types of animals (Fig. 5). The protein family analyses were performed for the 34 newly reconstructed (Table 2) and the 15 published Lak genomes, and the 181 circularized huge phage genomes reported recently ^16^. Clustering analyses grouped the ~660 kbp phage with other Lak phages (Supplementary Fig. S10), although it has 294 unique protein families (Supplementary Table 5). A total of 224 protein families were detected in at least 47 out of the 49 Lak genomes (referred to as “Lak_core”; Supplementary Table 5). Among “Lak_core’’ protein families, 114 were only present in Lak genomes (i.e., Lak-specific). Only 3 Lak-specific protein families could be annotated (i.e., magnesium transporter (K03284), protein transport protein SEC20 (K08497) and a pyruvyltransferase-like protein (K13665)). Interestingly, the pyruvyltransferase-like protein contains two domains (Glyco_tranf_2_4 and PS_pyruv_trans), both of which have the highest similarity to those from *Prevotella* spp. (Supplementary Fig. S11). A total of 110 “Lak_core” protein families are also present in non-Lak genomes. They generally are phage structural proteins, including large terminase, prohead core protein, baseplate wedge subunits, neck protein and tail tube protein etc., and those for replication, recombination and repair including HNH nucleases, DUTP diphosphatase, DNA polymerase, DNA primase, RecA/RadA recombinase and ribonucleoside-triphosphate reductase etc. (Supplementary Table 5).

**Fig. 5.**
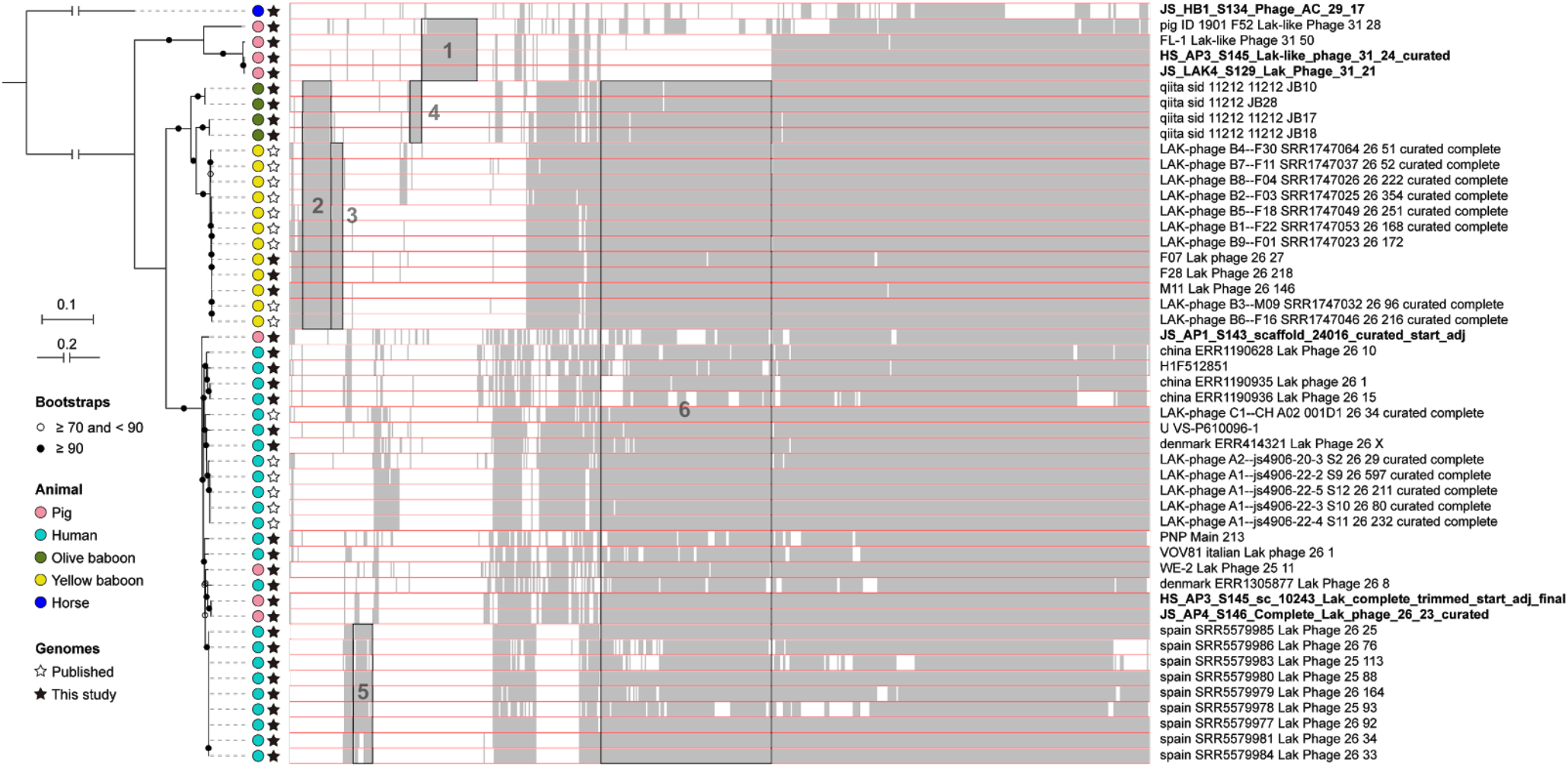
Phylogenomic analyses of the 49 Lak phage genomes. The phylogenetic tree (left) was built based on concatenated sequences of 49 single copy protein families detected in all Lak genomes and rerooted using the sequence of the ~660 kbp horse-associated Lak phage. The protein family content heatmap (right), aligned with the phylogenetic tree, shows the presence/absence of protein families that could be detected in at least 4 genomes. The names of the 6 complete Lak genomes reported in this study are in bold. A total of 6 blocks of protein families with animal-specific distribution patterns are highlighted in boxes and numbered: 1 = GC31 pig group specific, 2 = baboon specific, 3 = yellow baboon specific, 4 = olive baboon specific, 5 = Spanish human enriched, 6 = GC26 group enriched.

We detected some protein families in Lak genomes that are only found in specific animal hosts (Fig. 5). For example, 18 protein families were only detected in baboon Lak genomes, three only in olive baboon Lak genomes and 6 only in yellow baboon Lak genomes. Also, we found 37 protein families in all four genomes of the pig-associated GC31 group (including UK and Danish pigs) but in no other Lak genomes. We speculate that these animal host-specific protein families could be important during infection of their animal-specific *Prevotella* species and/or adaptation to the animal host. However, the inability to assign functions to these proteins at present hinders our understanding of their biological roles (Supplementary Table 5).

## Discussion

### Lak phages are prevalent across diverse human and animal microbiomes

The present study extends our previous finding of Lak phages in gut microbiomes of humans (Bangladesh and Hadza tribe), Danish pigs, yellow baboons and, in some cow rumens ^11^. Here, we show that Lak phages are present in microbiomes of additional human cohorts (China, Denmark, Italy, Spain, Israel and Liberia), various pig breeds, non-human primates (white-naped mangabey, yellow and olive baboons, macaques, guenons and colobus), horses, warthogs, fallow deer, a cow rumen, and Galapagos giant tortoises (first reported from a reptile) which likely had similar microbiome composition to hindgut-fermenting mammals (*Bacteroidetes* and *Firmicutes* dominated) ^19,20^. Lak was mainly detected in monogastric (single-chambered stomach) hindgut fermenters, but also in some ruminants (cow and deer). The Lak genome from one horse is notable because it is now the largest gut-associated phage genome (659,950 bp), and is one of the largest phage genomes reported to date ^16^. We are confident regarding this expanded genome size range, because key genomes (including the largest) were manually curated to completion. Overall, our findings demonstrate that diverse Lak phages are prevalent in microbiomes of humans and animals across the globe.

Phylogenies group together Lak phages that inhabit humans and some pigs, separating them from phages from other pigs (i.e., GC31 group), and from phages in non-human primates (and likely other animals based on PCR sequences). Different phylogenetic groups correlate with different protein family contents. Animal-specific protein families seem independent of geographic origins, but the lack of functional predictions precludes an explanation. The distribution of similar Lak genotypes may be influenced by animal physiology (e.g., contributing to similar phages amongst non-human primates), the potential for transmission (which could partly explain related Lak phages in humans and domestic pigs), and the distributions of potential bacterial host species or strains. It is possible that Lak phages migrate independent of their bacterial hosts and have a broad host range. Effects due directly to animal physiology are possible, but similarities in diet, and thus *Prevotella* species or strain composition of gut microbiomes, most likely influences the distribution of Lak phages.

### Distribution of Lak phage and *Prevotella* across pig gastrointestinal sites

Pigs were used in this study as a model system to analyse the distribution of *Prevotella* and Lak phages throughout the monogastric digestive tracts. It should be noted that Lak host range within the *Prevotella* genus is uncertain, so the broad *Prevotella* qPCR primers that were used ^21^ may not exclusively target the *Prevotella spp.* that the Lak phages infect. Nonetheless, results for the distribution of *Prevotella* and Lak phages align with the current knowledge that *Prevotella* are common in pigs, and are enriched in the hindgut (main site of fibre fermentation), whereas genera within the phyla *Firmicutes* and *Proteobacteria* dominate the foregut ^9,22–25^. Within the foregut and hindgut compartments, the absolute abundances of Lak and *Prevotella* in the lumen and mucosa do not differ at GIT sites (*P* < 0.05; Figs. 1b and S5), despite the fact that *Prevotella* are able to degrade mucins and are equipped to colonise the mucosa ^26^. Theoretically, the relative abundance of Lak phages compared to *Prevotella* might also be higher in the mucosa compared to the lumen, because adhesion of phages to the mucosa should increase phage-bacteria encounter rates ^27,28^. The finding that this is not the case may relate to the counteracting effect of the mucosa allowing bacteria to evade phage predation ^27,28^. Notably, the relative abundances of Lak compared to *Prevotella* were higher in the foregut compared to the hindgut mucosa (Fig 1c). This may be a consequence of slower digesta transit times through the hindgut compared to foregut lumens^29^, coinciding with increased establishment and thus higher relative abundance of *Prevotella* in the foregut mucosa compared to other bacteria ^9,22–25^. This may increase the probability of successful Lak phage replication in the foregut compared to the hindgut.

### Factors affecting the prevalence of Lak in pigs

Phenotypic differences in pig breeds, sex and age can affect microbiome composition ^10,30^. Lak-positive pigs represented a variety of breeds, ages and both sexes. However, Lak was most frequently detected in finisher pigs (*n* 13), and not detected in piglets or gestating sows. Colonisation of the piglet GIT is facilitated by the sow through birth and lactation ^31^, but their microbiomes are highly unstable and a *Bacteroides* to *Prevotella* shift often occurs as maturity is reached ^8,32^. Lak prevalence in finisher pigs could relate to dietary provision of fibrous ingredients being greater than in other production stages, but lower than in gestating sows (e.g. 17.5% wheatfeed + 5% rapemeal (Cambridge dry sow) vs. 5% wheatfeed + 7% rapemeal (Cambridge finisher); Supplementary Table 6). This may have increased microbial diversity and reduced the prevalence of *Bacteroidetes* in gestating sows compared to finisher pigs ^33,34^. Overall, dietary differences that reduced *Prevotella* relative abundances may explain the non-detection of Lak in piglets and gestating sows (Supplementary Table 6).

### Possible significance of *Prevotella* lysis

Lak phage predation could shape *Prevotella* population structure and overall microbiome composition. This is important because, although a commensal in various microbiomes, *Prevotella* has been linked to a variety of human diseases ^21,35–40^. *P. copri* overgrowth in the gut has been linked to rheumatoid arthritis in humans ^41–43^. *P. bivia* is strongly associated with bacterial vaginosis ^21,36,40^, and recently severe pre-eclampsia in humans^44^. We detected Lak phage in pig vaginal mucosas, albeit at lower abundance than in rectums (Supplementary Table 3). *P. copri* and *P. bivia* are common in both pigs ^45^ and humans ^45,46^, thus it is possible that Lak predation of these bacteria could reduce the incidence of their associated diseases.

In humans and animals, *Prevotella* lysis by Lak phages may affect fibre fermentation, with potential health implications. In pigs, *Prevotella*-dominated enterotypes are associated with improved growth performance ^8,9^. Given that *Prevotella* are enriched in the hindgut where fibre is primarily fermented, lysis could be detrimental to the animal host. However, overgrowth of certain *Prevotella* species in pigs may reduce feed efficiency, and facilitate undesirable fat accumulation ^47–49^. Thus, Lak phage predation could positively or negatively impact swine production. Besides the presence of a caecum, the pig gut physiology and microbiome composition is comparable to humans ^10^. Therefore, the distribution of Lak phage and *Prevotella* in the swine GIT could inform our understanding of Lak and *Prevotella* distributions more generally.

## Conclusions

Lak are substantially more diverse and have a wider range of genome sizes and genome GC contents than realised previously. All lineages appear to use the same alternative genetic code. Lak phages occur in the microbiomes of many humans and animals including reptiles with the largest detected in a racehorse; this is the largest gut phage reported to date. Lak appear to be particularly common in pig microbiomes, where they are found in multiple body sites and enriched in the hindgut. It may be possible to harness Lak phages to modulate microbiome structure and composition, with long term implications for treatment of human diseases, including rheumatoid arthritis and vaginosis, and to improve swine growth performance.

## Author contributions

The study was designed by J.M.S. and J.F.B.. M.A.C. extracted DNA from samples, designed Lak PCR and qPCR primers, performed PCR amplification, qPCR (with support from P.C.). J.F.B., L.X.C., A.D., A.X., A.T., and N.S. performed metagenomic assembly and initial genome binning of Lak phages. A.E.D., R.S., A.S. and S.L. handled metagenomic data and assembly. J.F.B. and L.X.C. performed manual genome curations. L.X.C. performed phylogenetic analyses and protein family analyses. A.L.B. performed the tRNA analyses and contributed to analysis of alternative coding. F.D. coordinated post-mortem pig sampling, removed entire guts and lungs, and collated info from producers. M.B. advised on post-mortem pig sampling, removed gut sections for dissection. M.A.C and R.B. dissected mucosal tissues and digesta from pig guts. F.D. and R.B. dissected pig lung and vaginal tissues. R.M.W. and M.A.H. provided Cambridge pig faecal samples and meta-data. L.R. coordinated zoo animal sampling. N.B. performed the phylogenomic analyses. M.A.C, L.X.C. and J.F.B. wrote the manuscript with input from J.M.S., A.L.B. and N.B. All authors contributed to editing the manuscript.

## Acknowledgments

Funding for this study was provided by the Innovative Genomics Institute, UC Berkeley, the Chan Zuckerberg Biohub, and the National Institutes of Health (RAI092531A) to J.F.B. M.A.C. was funded by the BBSRC (BB/M009513/1) through the London Interdisciplinary Doctoral Program. N.B acknowledges funding by the BBSRC (BB/R009597/1). M.A.H. acknowledges funding from the MRC (MR/N002660/1) and R.M.W. acknowledges funding from the BBSRC (BB/M011194/1) for their contributions. A.L.B was supported by a Miller Basic Science Research Fellowship at UC-Berkeley. N.S. was supported by the European Research Council (ERC-STG project MetaPG-716575). For their contributions to this study, we thank Christopher Davies and Steven Van Winden (RVC), Harriet Barlett (University of Cambridge), Roger and Sharron Ingram (Wendover Stables, Epsom), Caroline Rymer and David Humphries (University of Reading), Katherine Thompson (Birkbeck, University of London), Andrew Mead and Ludovic Pelligand (RVC), Mehmet Davrandi and Sean Nair (Eastman Dental Institute, UCL), and Jennifer Safapour.

## Methods

### Animals and sampling

A total of 187 samples from different animals were screened by PCR (Supplementary Table 2). In addition to faecal samples, digesta and mucosal tissues were obtained where possible. At Langford Abattoir (University of Bristol, UK), finisher pigs (*Sus scrofa*) were fasted for 24-hours before arrival, where they were stunned and humanely slaughtered for gut, vaginal and lung sampling. Pigs 1-4 and 7-11 (Welsh x Petrain) came from a different smallholding to pigs 5-6 (Welsh). For GIT sampling, pigs were reared in pairs with 1 male and 1 female (1-2, 3-4, 5-6), and each pair was reared separately. Vaginal, lung and rectal samples from female pigs 7-11 were harvested separately. Pig faeces from a commercial farm was obtained and supplied by The University of Cambridge, UK (Large white x Landrace x Hampshire). Cambridge samples pertained to various production stages: 2 piglets, 2 pre-farrow sows, 3 early-gestation sows, 1 late-gestation sow, 5 weaner pigs (8-12 weeks), 2 grower pigs (12-18 weeks), 2 finisher pigs (18-22 weeks).

Rumen-cannulated dairy cows (*Bos taurus*, Holstein) were also sampled (Centre for Dairy Research, CEDAR, University of Reading, UK). Frozen ROSS 308 broiler (*Gallus gallus domesticus*) caecal digesta was obtained from a feeding trial at The Royal Veterinary College (RVC, UK). Only samples from untreated, control birds were used. Available animal diet composition is listed in Supplementary Table 6.

### Sample collection

To avoid cross-contamination, gloves were changed between each sample and only sterile equipment and collection tubes were used. For all faecal samples, approximately 2 g faeces was taken from the centre of the sample to limit environmental contaminants. Dairy cows at CEDAR were moved to individual pens and cannulas opened for rumen fluid collection. Rectal samples were also taken from the same 3 cows. Cambridge pig samples were collected in sterile 7 ml tubes and frozen at −80°C, before transfer to The Santini Lab, University College London (UCL, UK) on dry ice. All other faeces, cow and broiler digesta were transferred on ice packs to UCL within 3 hours and stored at −80°C until analysis.

At Langford abattoir, (after scalding) entire GIT’s from 6 post-mortem finisher pigs (1-6) were removed from oesophagus to rectum, within 30 minutes of slaughter. Digestive compartments were sectioned with cable ties and removed: mid-jejunum, terminal ileum (10 cm anterior to ileo-caecal junction), proximal spiral (10 cm distal to ileocecal junction), distal spiral, distal caecum and rectum. Luminal digesta and mucosal scrapings were collected using ethanol-sterilized equipment. Ileal lumens were empty in 2/6 pigs. Vaginal and rectal samples from pigs 7-11 were also obtained before scalding, vulvas were sanitised with 100% ethanol and vaginal mucosas were removed using sterile equipment, rectal samples were then collected using clean spatulas. Lung sampling from the same animals was carried out post-scalding; tracheas were clamped to avoid scalding lung contents before longitudinal dissection of each lung following the left and right bronchi. Mucosal scrapings of each lung were taken with sterile scalpels and pooled for each pig. All post-mortem samples were flash-frozen and transported on dry ice to UCL and stored at −80°C.

### DNA extraction

All samples were thawed at room temperature and DNA was extracted using a QIAamp PowerFecal DNA kit (Qiagen, Hilden, Germany), following the manufacturer instructions. DNA concentration and 260/280 Ratio were measured in duplicate using NanoDrop™ 2000 (ThermoFisher Scientific, MA, USA) and averaged, to ensure sufficient DNA quality and concentration for PCR.

### PCR and amplicon sequencing

The genes for major capsid protein (MCP), portal vertex protein (PVP) and tail sheath monomer (TSM) from human and baboon Lak genomes were aligned using ClustalW in MEGA-10 ^50^ to identify homologous regions. Primers were designed in Primer-BLAST^51^ and synthesized by Sigma-Aldrich (MO, USA). The designed primer pairs were specificity checked and optimised; LakMC581-F: 5’-GGAGTCATACGAACACCAGAAGT-3’ / LakMC1053-R: 5’-GTAGTTCTTACACTTCACGCTCCTC-3’ (MCP amplicon: 473 bp), LakPV767-F: 5’-CATGGTCAACAGGTATGTATGG-3’ / LakPV1261-R: 5’-CCTCTCGTGTTATACTTGCATCA-3’ (PVP amplicon: 495 bp), LakTS3039-F 5’-CTTCCATCTAAGAGACAGTTTGA-3’ / LakTS3781-R: 5’-GCTATGATGTCCGGTGTGTTG-3’ (TSM amplicon: 689 bp). Each 25 μL PCR reaction contained 150 ng/μL template DNA (alongside a swine positive control), 5.5 μL master mix, free deoxynucleotides (dNTPs, 200 μM), forward and reverse primers (0.14 μM), NH_4_ reaction buffer (1 x), and MgCl2 (3 mM), and 1.25 U BIOTAQ™ DNA Polymerase (Bioline, London, UK). A Mastercycler Nexus GSX1 (Eppendorf, Germany) was programmed for 40 cycles with DNA denaturation temperature of 96°C for 10 s, annealing (MCP: 61°C, PVP: 58°C, TSM: 57°C) for 30 s, and extension at 72°C (MCP and PVP: 15s, TSM: 20s) with a final extension of 5 min. PCR amplicons were visualised by agarose gel electrophoresis. PCR products were purified using either a QIAquick PCR purification or gel extraction kit (Qiagen, Hilden, Germany). Sanger sequencing of purified PCR products was performed by Eurofins, Germany. BLASTN^52^ was used to confirm sequences were similar to Lak. Forward and reverse sequences were aligned using MEGA-X^50^, and quality checked against sequence chromatograms. Three genes (Lak MCP, TSM and PVP) were sequenced for all animal cohorts, except giant tortoise (GTA), fallow deer (FD) and pig 2 jejunal mucosa (JM2), where two of the three genes were sequenced. A summary of sequences obtained for each sample is reported in Supplementary Table 1.

### Quantitative real-time PCR (qPCR)

Lak phage and *Prevotella* abundances were determined by quantitative PCR (qPCR) using the standard curve method. *Prevotella* genus-specific 16s rRNA primers designed previously for the human vaginal microbiome were used ^21^. Primer-BLAST^51^ was used to check coverage for common gut *Prevotella* species. These included strains of *P. copri, P. stercorea, P. melaninogenica, P. intermedia, P. jejuni, P. bivia* and *P. nigrescens,* many of which are found in pigs and humans ^45,46^. The Lak MCP genes from available pig metagenomes ^11^ were aligned by ClustalW in MEGA-X ^50^. Lak candidate primers with amplicons 114-221 bp were designed in Primer-BLAST^51^ and synthesised by Sigma-Aldrich (MO, USA), along with *Prevotella* primers. Primer pairs were checked for primer dimers and hairpins in OligoAnalyzer (Integrated DNA Technologies Inc., Iowa, USA) and specificity-checked by Sanger sequencing PCR products prior to use in qPCR (Eurofins, Germany).

LakMC581-F/LakMC1053-R PCR product from pig rectal DNA was used to generate Lak standards, as this encompassed qPCR targets. *P. copri* DNA (DSM 18205, type strain) was used for *Prevotella* standards. Serial dilutions (9 x 1:10) starting at 5 ng DNA were used for standard curves (quantification cycle (Cq) vs. Log DNA dilution) during quantification and to determine primer efficiencies ((−1+10^-1/slope^) x 100). The selected Lak primer pair yielded an efficiency of 102.8%, LakMCP683-F: 5’-CAACCAAGAGCGAACAAACGAG-3’ / LakMC803-R: 5’-TAACAGACCTTCAGAAACAGTGGG-3’ (amplicon: 121 bp). The *Prevotella* primer pair^21^ yielded an efficiency of 94.1%, Prevo-F: 5’-GGGATGCGTCTGATTAGCTTGTT-3’ /Prevo-R: 5’-CTGCACGCTACTTGGCTGGTTC-3’ (amplicon: 179 bp).

Data were obtained using a PikoReal™ real-time PCR system (Thermo Fisher Scientific, MA, USA), with a QuantiNova SYBR Green PCR kit (Qiagen, Germany). 9 uL master mix providing 1x SYBR Green master mix, 0.7 uM primers, 1x ROX passive reference dye and 1 uL nuclease-free water, was pipetted into Piko 96-well plates (Thermo Fisher Scientific, MA, USA). 1 uL gDNA, diluted in 1 x template dilution buffer, was added to the master mix (10 uL reaction volume). Plates were sealed using Piko Optical Heat Seals (Thermo Fisher Scientific, MA, USA). 3 technical replicates and no template controls (NTC) were included throughout. Standards were run in parallel to 10 ng sample microbiome DNA. Primer efficiencies remained at 90-103 %, and melt curves suggested no nonspecific binding or secondary structures. Representative standard and melt curves are shown in Supplementary Figs. S2 and 3. Lak and *Prevotella* quantities (ng) were extrapolated from standard curves and collated into a database.

Copy numbers were calculated, log-transformed, and technical replicates averaged. The qPCR data were analysed in JMP^®^ Pro 14.1 (SAS Institute Inc., NC, USA, 2019). Distribution was analysed by GIT site for both mucosal and lumen log copy numbers (Supplementary Fig. S5). No outliers were identified 1.5*IQR. A Shapiro Wilk-Test for normality suggested data were near normally distributed (*P* > 0.05). To compare Lak phage abundances, standard least square mean comparisons were made using a full factorial approach and restricted maximum likelihood (REML) method. ‘GIT site*Sample type’, ‘Sex’ and ‘Farm’ were included as fixed effects, and plate number as a random effect, to account for co-variation. Actual vs. predicted values indicated adequate model fit (R^2^=0.96, RMSE=0.36, *P* < 0.0001). The abundance of Lak phage copies / *Prevotella* copies were calculated to estimate phage copies per host genome; a fixed effect model was used with the same parameters, but plate number was ommited. Treatment means were separated using Tukey’s HSD test (α = 0.05 and 0.001). Least square comparisons were made between vagina and rectal samples with no covariates (as these animals were of the same sex, from the same farm, and qPCR was performed on a single plate). Raw qPCR data and statistical outputs are included in Supplementary Tables 7 and 8.

### Metagenomic sequencing and analyses

A total of 31 samples confirmed with Lak phages were sequenced. The raw reads of each metagenomic sample were filtered to remove Illumina adapters, PhiX and other contaminants with BBTools ^53^, and low-quality bases and reads using Sickle (version 1.33, https.github.com/najoshi/sickle). The high-quality reads of each sample were assembled using idba_ud ^54^ (parameters: --mink 20 --maxk 140 --step 20 --pre_correction), or MEGAHIT ^55^ (parameters: --k-list 21,29,39,59,79,99,119,141). For a given sample, the high-quality reads of all samples from the same sampling site were individually mapped to the assembled scaffold set of each sample using Bowtie 2 with default parameters ^56^. The coverage of a given scaffold was calculated as the total number of bases mapped to it divided by its length. The scaffolds with a minimum length of 1 kbp were uploaded to the ggKbase platform. The protein-coding genes were predicted using Prodigal ^57^ (-m -p meta) from scaffolds and annotated using usearch ^58^ against KEGG ^59^, UniRef ^60^ and UniProt ^61^. Some published metagenomic datasets (Supplementary Table 1) ^15,62^ were also analysed using the same pipeline as described above..

### Manual genome curation

The *de novo* assembled contigs/scaffolds were searched against the 15 published Lak genomes ^11^ using BLASTN^52^. The BLAST hits were filtered to retain those with an alignment longer than 2 kbp and a minimum similarity of 90%. The target contigs/scaffolds from a given sample were grouped into bin(s) based on their GC content and coverage. Manual genome curation was performed on the bin(s) as previously described ^63^ by read mapping, scaffold extension and join, and manual fixation of assembly errors, attempt for completion was also conducted until a circularized genome was obtained.

### CRISPR-Cas analyses

All the predicted proteins of scaffolds with a minimum length of 1 kbp were searched against local HMM databases including all reported Cas proteins, and the nucleotide sequences of the same set of scaffolds were scanned for CRISPR loci using minced ^64^ (-minSL = 17). The spacers were extracted from the scaffolds with CRISPR loci as determined by minced, and also from reads mapped to these corresponding scaffolds using a local python script as previously described ^65^. For the published genomes, only spacers from the scaffold consensus sequences were extracted, as no mapped reads are available. Duplicated spacers were removed using cd-hit-est (-c = 1, -aS = 1, -aL = 1) and the unique spacer sequences were used to build a database for BLASTN ^52^ searches (task = blastn-short, e-value = 1e-3) against the Lak genomic sequences. Once a spacer was found to target a Lak phage scaffold with at least 90% alignment similarity, the original scaffold of the spacer was checked for a CRISPR locus and Cas proteins.

### tRNA analysis

The tRNAs were predicted using tRNAscan-SE ^18^ in eukaryotic mode with default settings. Lak tRNAs have been previously established to contain introns and thus are not all classified in bacterial mode.

### Phage protein family analyses

Protein family analyses were performed as previously described ^66^. In detail, first, all-vs-all searches were performed using MMseqs2 ^67^, with parameters set as e-value = 0.001, sensitivity = 7.5 and cover = 0.5. Second, a sequence similarity network was built based on the pairwise similarities, then the greedy set cover algorithm from MMseqs2 was performed to define protein subclusters (i.e., protein subfamilies). Third, in order to test for distant homology, we grouped subfamilies into protein families using an HMM-HMM comparison procedure as follows.

The proteins of each subfamily with at least two protein members were aligned using the result2msa parameter of MMseqs2, and HMM profiles were built from the multiple sequence alignment using the HHpred suite ^68^. The subfamilies were then compared to each other using hhblits ^69^ from the HHpred suite (with parameters -v 0 -p 50 -z 4 -Z 32000 -B 0 -b 0). For subfamilies with probability scores of ≥ 95% and coverage ≥ 0.5, a similarity score (probability × coverage) was used as the weights of the input network in the final clustering using the Markov CLustering algorithm ^70^, with 2.0 as the inflation parameter. Finally, the resulting clusters were defined as protein families. The clustering analyses of the presence and absence of protein families detected in the phage genomes were performed with Jaccard distance and complete linkage.

### Phylogenetic analyses

To reveal the phylogeny of Lak phages reconstructed in this study. The shared single copy gene product sequences from each genome were concatenated and aligned with MAFFT (default parameters) ^71^. The alignment was subsequently converted into a phylogenetic tree on the MAFFT web-server using 100 bootstraps, Neighbor Joining, JTT as a substitution model ^72^ and visualized in iTOL ^73^.

## Data availability

The 34 newly reconstructed Lak megaphage genomes have been deposited at NCBI under BioProject PRJNA688310, and also available from ggkbase https://ggkbase.berkeley.edu/Lak2/organisms (please sign in by providing your email address to download) and at figshare (https://figshare.com/articles/dataset/34_new_Lak_phage_genomes/13493721). The NCBI accession information for all published datasets is available from Supplementary Table 1.

