## Supplementary Figures for "Wide distribution of alternatively coded Lak megaphages in animal microbiomes"

### Study design and main results

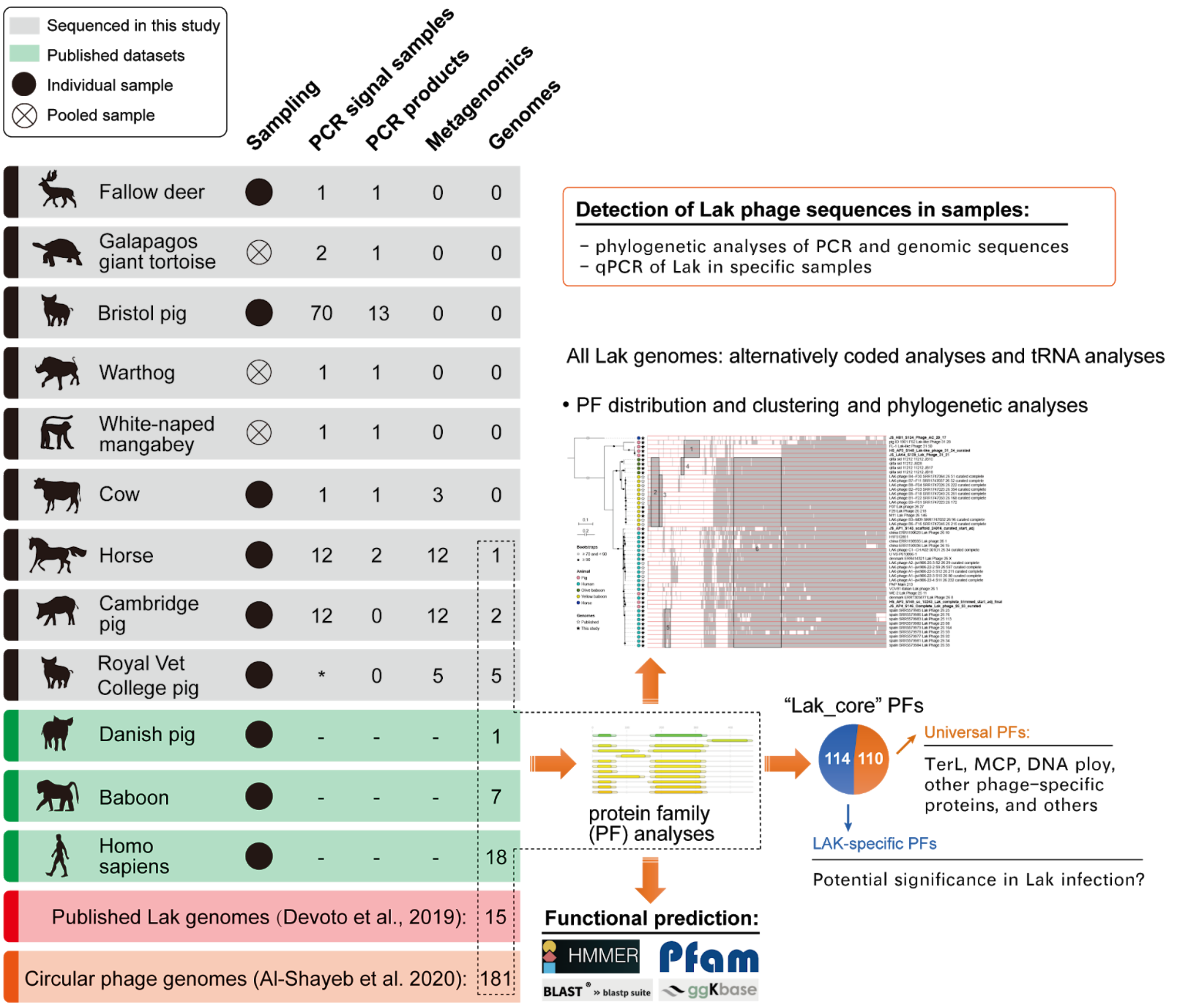


**Supplementary Fig. S1 | Graphical abstract showing the study design, methods and main results.** Sanger sequences from Lak major capsid, tail sheath monomer and portal vertex PCR amplicons were included for animal cohorts where genomes have not been resolved. Cambridge pig sequences represented a weaner, grower and finisher. Bristol pig sequences represented jejunal, ileal, proximal spiral, distal spiral, caecal and rectal lumen contents from one pig, along with rectal contents from another pig; these samples were also subjected to qPCR analyses. 3 genes were sequenced for all cohorts except Fallow Deer and Bristol Pig 2 jejunal mucosa, where 2 genes were sequenced. * PCR experiment was not performed.

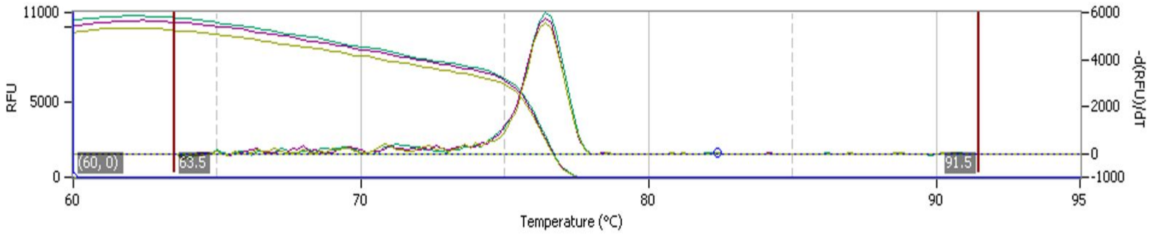


**a**

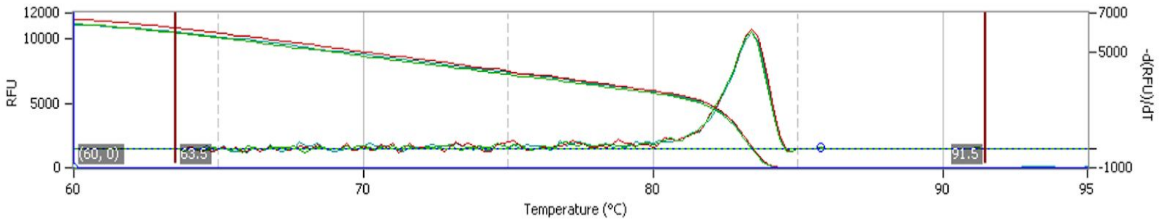


**b**

**Supplementary Fig. S2 | Representative melt curves for selected qPCR primer pairs.** RFU=relative fluorescence units using SYBR green. Single peak generated from 3 technical replicates indicates no nonspecific binding or secondary structures. (**a**) Representative result shown for Lak major capsid gene primers designed in the present study, using 10 ng Pig 2 Proximal Spiral (PS2) pooled digesta DNA. (**b**) Representative result shown for *Prevotella* genus-specific primers designed previously[17] using 10 ng Pig 2 Rectal (R2) pooled digesta DNA.

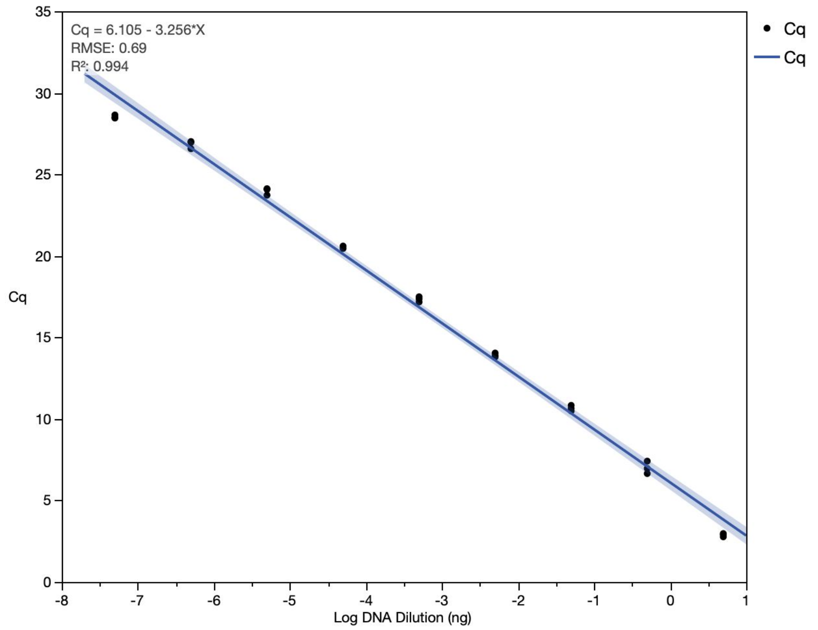


**a**

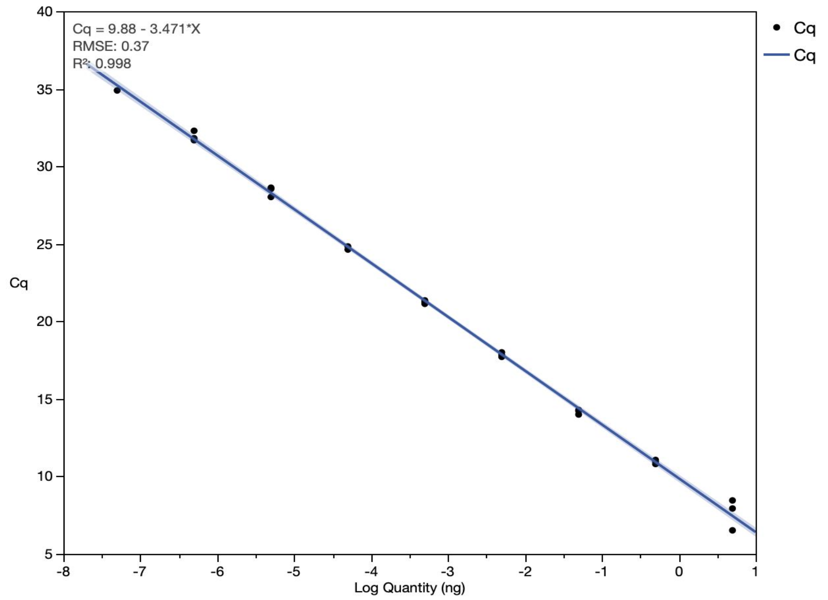


**b**

**Supplementary Fig. S3 | Standard curve for selected Lak qPCR primer pair.** Cq=Cycle of quantification; whereby SYBR green fluorescence increases above background signals; $R^2$=Coefficient of determination; RMSE=Root mean square error. Black dots represent technical replicates for each serial dilution: 1:10 starting at 5ng. Primer efficiencies (E) calculated from slopes ($(-1+10^{-1/\mathbf{-3.256}}) \times 100$): (**a**) Lak major capsid PCR amplicons used as standards with primers designed in this study, E = 102.8%; (**b**) *Prevotella copri* DNA (DSM 18205, type strain) used as standards with previously designed primers[17], E = 94.1%.

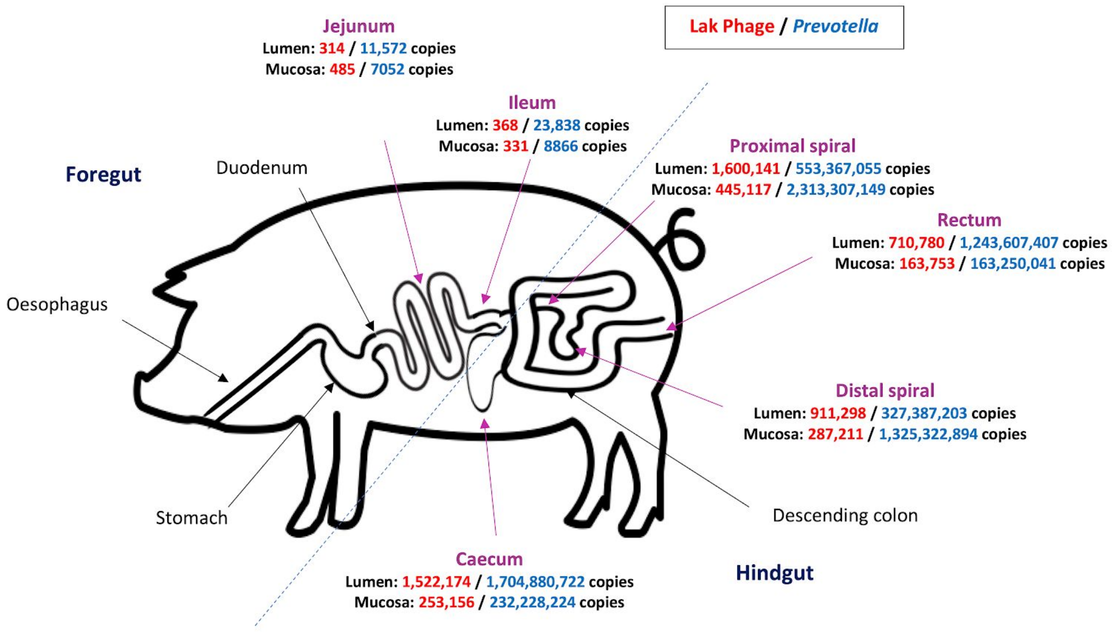


**Supplementary Fig. S4 | Abundance of Lak phage and *Prevotella* across the swine gastrointestinal tracts.** Pink arrows = gastrointestinal sites where Lak major capsid, portal vertex and tail sheath monomer genes were detected in digesta and mucosa by PCR. Black arrows = sites not sampled. Mucosa and Lumen copies = anti-log mean Lak major capsid and *Prevotella* 16s rRNA gene copy number, per 10 ng microbiome gDNA; determined by qPCR. Genes amplified are single copies and therefore = genome copies. Data represents 6 finisher pigs for all sites except ileal lumens; where digesta was only present in 4/6 pigs. Schematic not to scale.

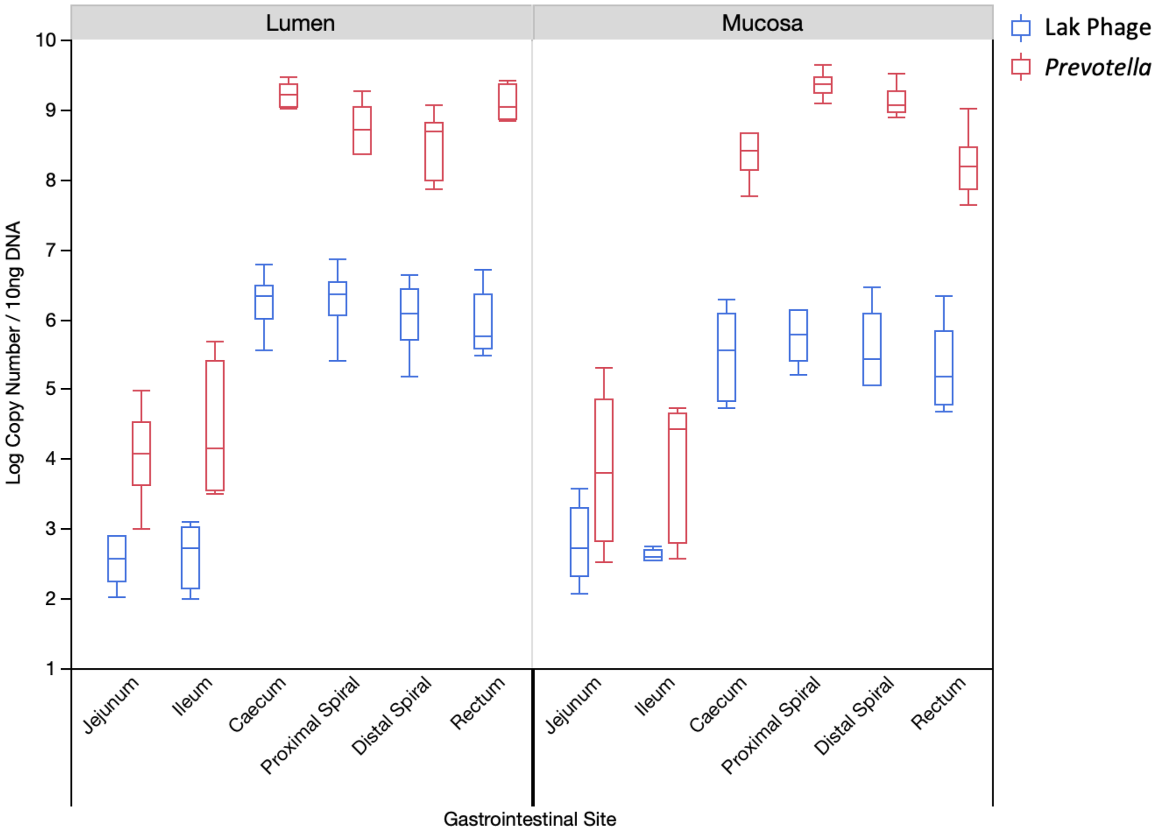


**Supplementary Fig. S5 | Distribution Lak phage and *Prevotella* log quantities across the swine gastrointestinal tract.** Top and bottom whiskers=minimum and maximum values. Box width=Interquartile range (IQR). Middle line=median. No outliers were identified 1.5*IQR. Shapiro-Wilk test for normality suggested log-transformed data were near-normally distributed. Data represents 6 finisher pigs for all sites except ileal lumens; where digesta was only present in 4/6 pigs.

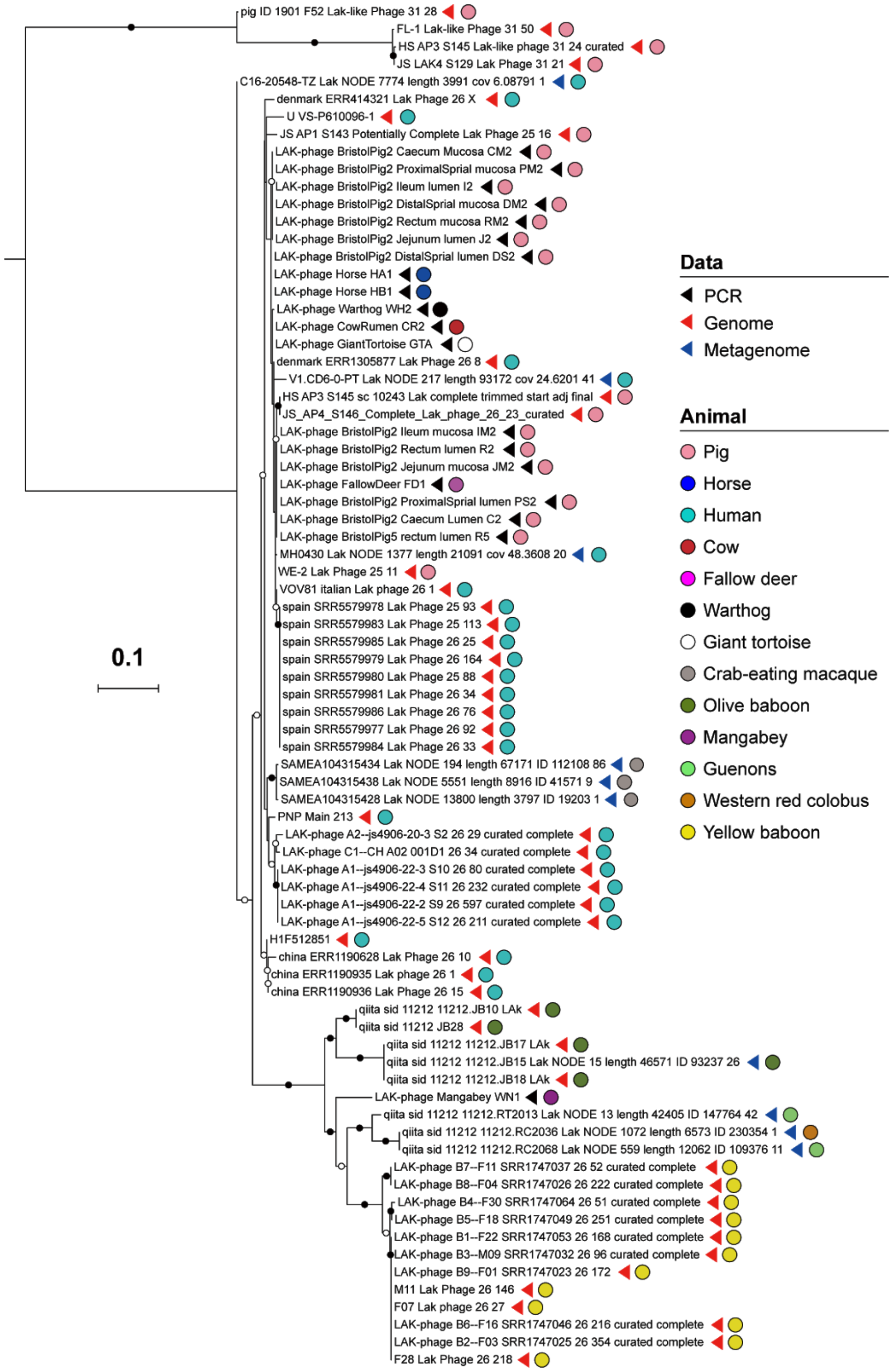


**Supplementary Fig. S6 |Phylogenetic analyses of Lak based on the sequences from PCR, genomes and metagenomes.** The nucleotide sequences encoding the portal vertex protein were aligned, and trimmed based on the length of the PCR sequences. The capsid of the 660 kbp phage is very divergent from others, thus excluded from the tree to have a better resolution of others. Bristol pig sequences obtained from the vaginal mucosa were identical to those found in the digestive tract.

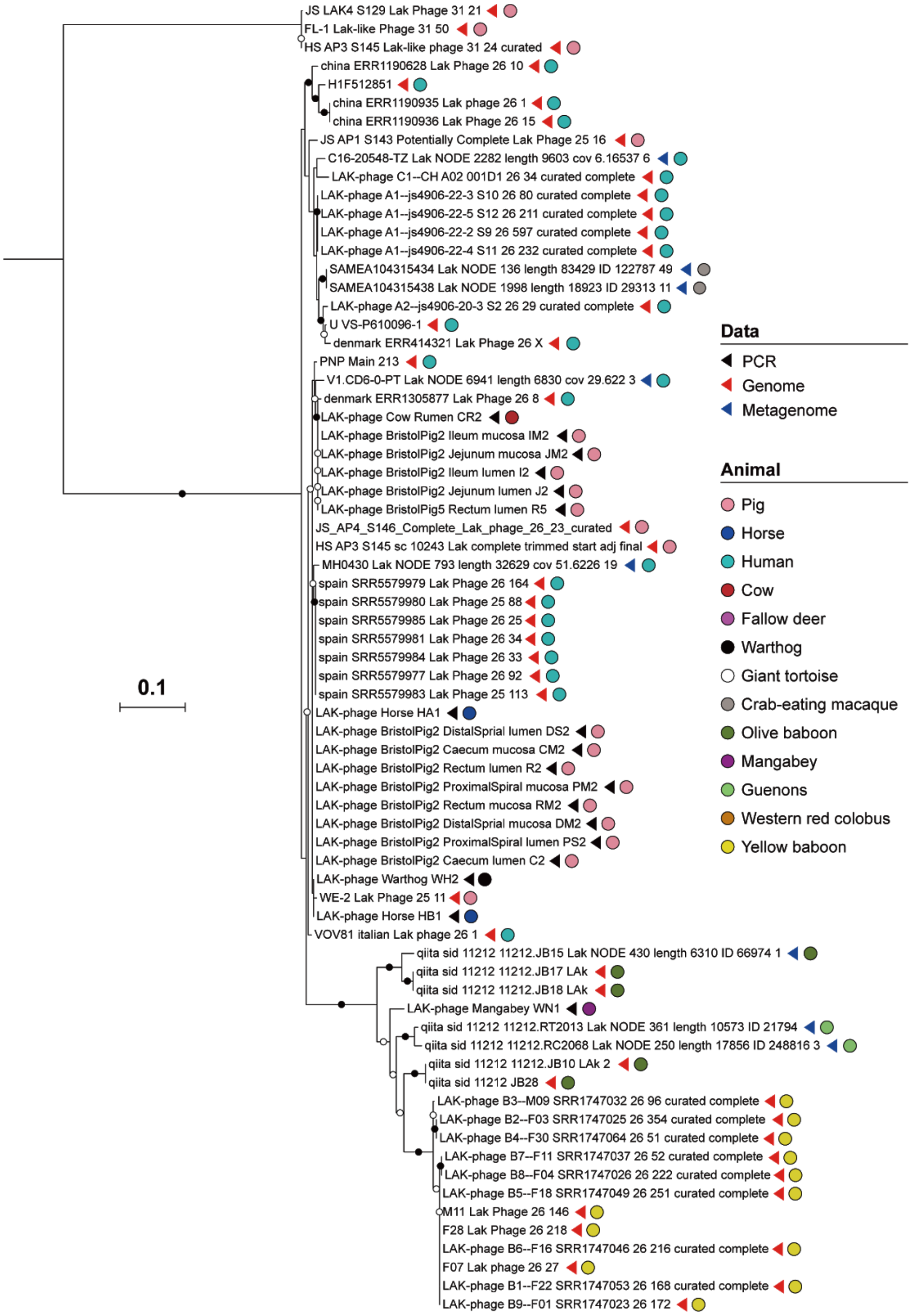


**Supplementary Fig. S7 | Phylogenetic analyses of Lak based on the sequences from PCR, genomes and metagenomes.** The nucleotide sequences encoding the tail sheath monomer were aligned, and trimmed based on the length of the PCR sequences. The capsid of the 660 kbp phage is very divergent from others, thus excluded from the tree to have a better resolution of others. Bristol pig sequences obtained from the vaginal mucosa were identical to those found in the digestive tract.

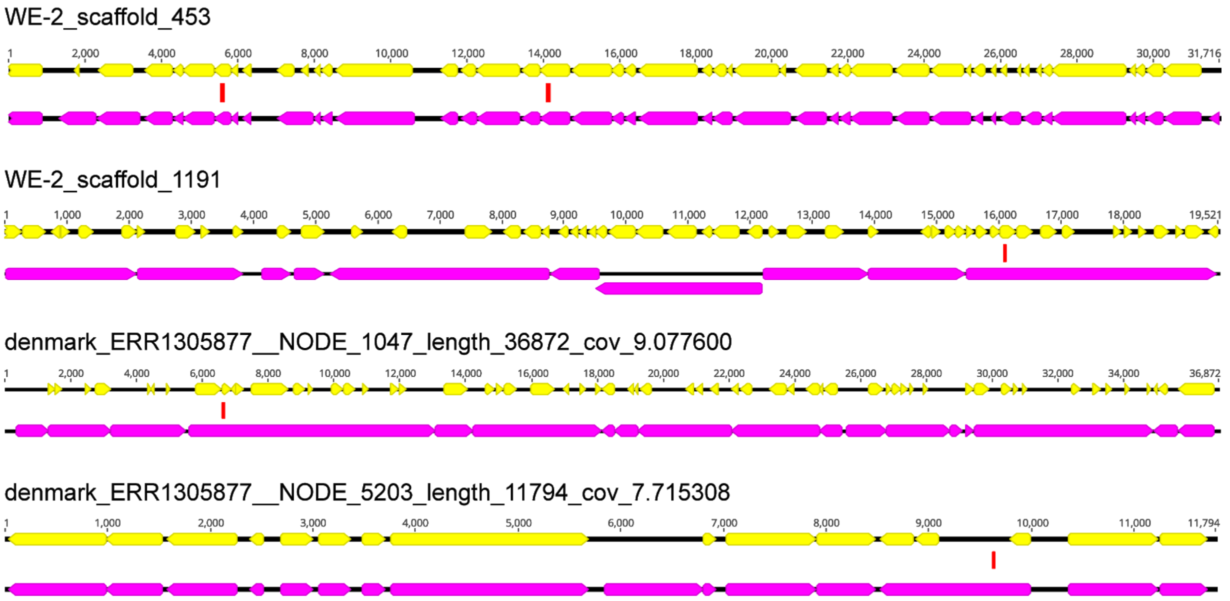


**Supplementary Fig. S8 | Examples of Lak phage sequences targeted by CRISPR-Cas spacers from *Prevotella* spp.** The targeted fragments by spacers are shown by red bars. For each scaffold, the protein-coding genes predicted with code 11 are shown in yellow, and those predicted with code 15 are shown in pink.

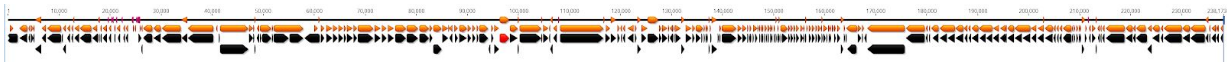

**Supplementary Fig. S9 | A 238 kbp phage genome fragment from a lineage divergent from Lak for which gene predictions using code 11 (black) and code 15 (orange) are essentially indistinguishable**. When tested, the novelty of a sequence precluded selection of a single vs. pair of open reading frames as most plausible.

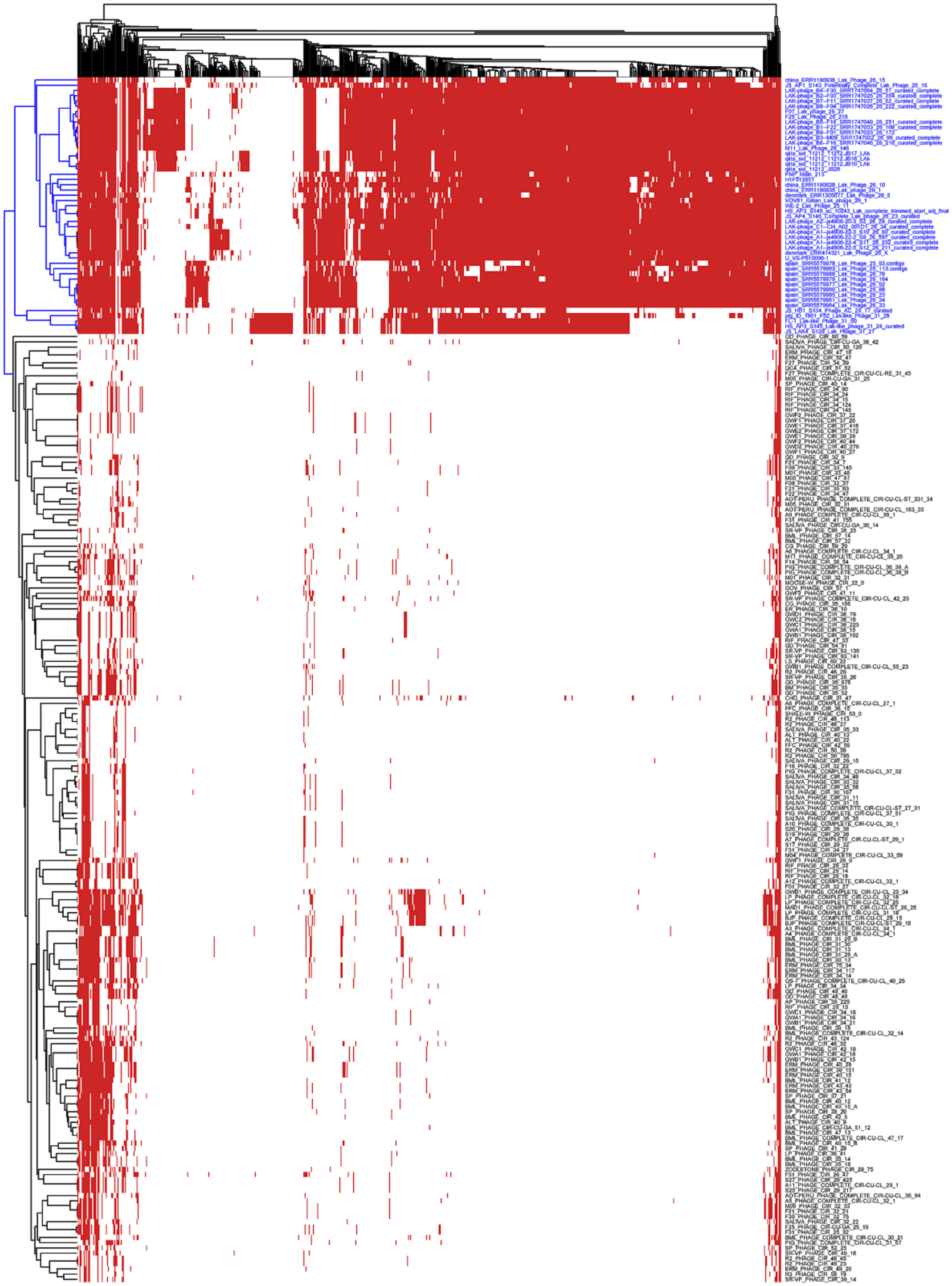

**Supplementary Fig. S10 | The clustering analyses of 49 Lak phages (including 15 published ones and 34 new ones reported in this study) and 181 huge phages reported with circular genomes based on the presence and absence of protein families.** Only those protein families detected in four or more Lak phage genomes were included. The clustering analyses were performed with Jaccard's distance and complete linkage.

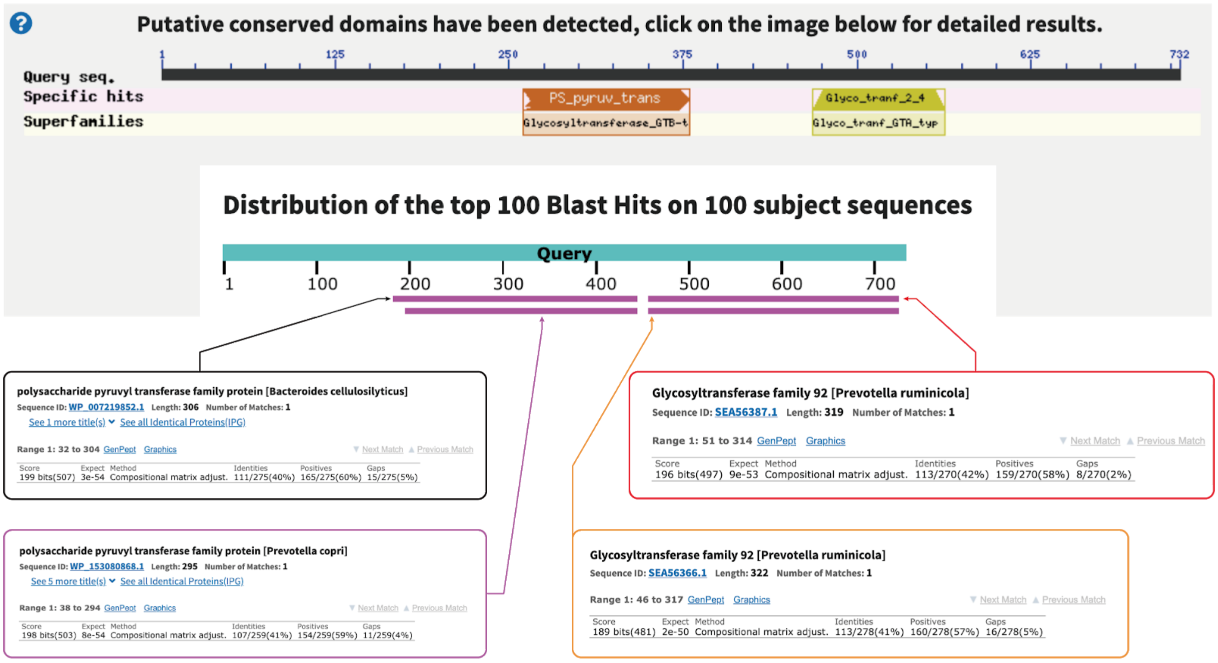


**Supplementary Fig. S11 | The sequence of a pyruvyltransferase-like protein from Lak phage is similar to that of *Prevotella* species.**
